# Positive-feedback defines the timing and robustness of angiogenesis

**DOI:** 10.1101/374389

**Authors:** Donna J. Page, Raphael Thuret, Lakshmi Venkatraman, Tokiharu Takahashi, Katie Bentley, Shane P. Herbert

**Author notes:** Correspondence: (K.B.), (S.P.H.).

## Abstract

Blood vessel formation by angiogenesis is critical for tissue development, homeostasis and repair, and is frequently dysregulated in disease[1–3]. Angiogenesis is triggered by vascular endothelial growth factor receptor-2/3 (VEGFR) signalling, which induces motile endothelial cell (ECs) tip identity[4,5]. Tip cells lead new branching vessels, but also coordinate collective EC movement by repressing tip identity in adjacent ECs via Delta-Like 4 (DLL4)-Notch-mediated down-regulation of VEGFR activity[6–13]. Hence, angiogenesis is driven by lateral inhibition-mediated competition of ECs for migratory status. Recent work reveals that temporal modulation of this DLL4-Notch-mediated lateral inhibition circuit fundamentally shapes both normal and pathological angiogenesis[14–17]. However, the core regulatory network defining the timing and dynamics of EC decision-making is unclear. Here, by integrating computational modeling with *in-vivo* experimentation, we uncover a unique ultrasensitive switch that temporally defines EC lateral inhibition and ultimately determines the timing, magnitude and robustness of angiogenic responses. We reveal that positive-feedback to Vegfr via the atypical tetraspanin, *tm4sf18*, amplifies Vegfr activity and expedites EC decision-making *in-vivo*. Moreover, this Tm4sf18-mediated positive-feedback confers robustness to angiogenesis against changeable environmental conditions by invoking bistable-like behavior. Consequently, mutation of *tm4sf18* in zebrafish delays motile EC selection, generates hypoplastic vessels, sensitizes ECs to fluctuations in pro-angiogenic signal and disrupts angiogenesis. We propose that positive-feedback transforms the normally protracted process of lateral inhibition into a quick, adaptive and robust decision-making mechanism, suggestive of a general framework for temporal modulation of cell fate specification in development and disease.

## RESULTS AND DISCUSSION

### Biphasic temporal control of angiogenesis by the Vegfr-Notch axis

To ensure appropriate angiogenic responses, excessive induction of tip identity is repressed by DLL4-Notch-mediated lateral inhibition[6–13]. During this process, VEGFR activation promotes up-regulation of the Notch ligand DLL4 in emerging tip cells, which *trans*-activates Notch in neighboring cells. Elevated Notch activity promotes down-regulation of VEGFR-2/3 function, rendering laterally-inhibited cells less responsive to VEGF signal (Figure 1A)[6–13]. As such, DLL4-Notch coordinates collective movement of sprouting EC populations, and loss of Notch results in EC hyper-sprouting *in-vivo*. However, we have little-to-no understanding of the underlying temporal features of these decision-making processes. Indeed, lateral inhibition is considered relatively slow, taking upwards of 6 h to complete multiple cycles of gene expression needed to amplify initially small differences in input signal[14,16–19], which is seemingly incompatible with the rapid dynamic changes in EC state, identity and behavior observed in angiogenesis[20,21].

**Figure 1.**
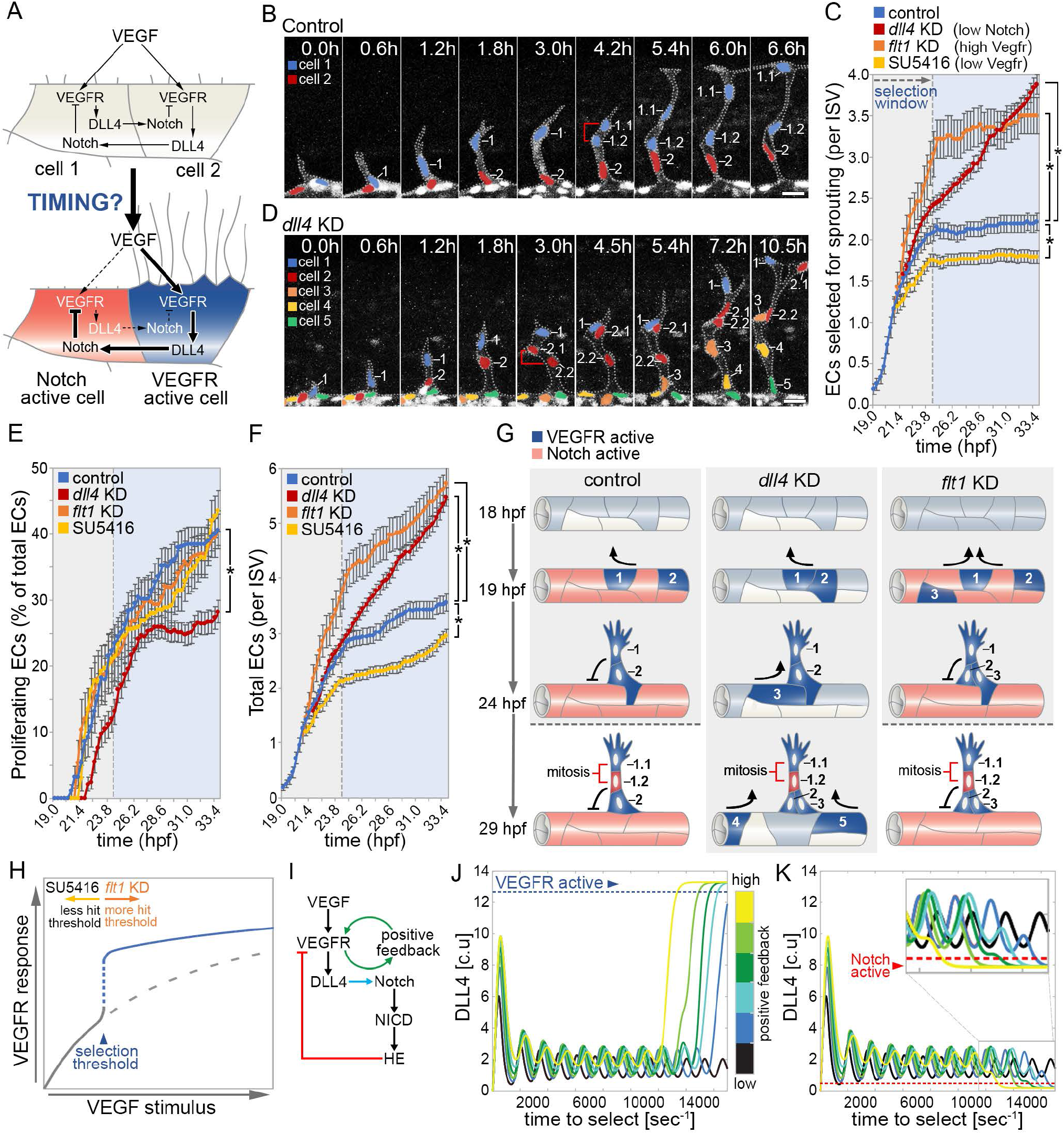
Temporal control of angiogenesis by Vegfr-Notch and positive-feedback. (A) Illustration of the VEGFR-DLL4-Notch lateral inhibition feedback loop between two ECs competing for tip selection. Following lateral inhibition, VEGFR activity increases in cell 2 whereas Notch activity increases in cell 1. (B-F) Time-lapse images of sprouting ISVs in control (B) and *dll4* knockdown (D) *Tg(kdrl:nlsEGFP)^zf109^* embryos from around 19 hours post-fertilisation (hpf). Brackets indicate dividing cells. Nuclei are pseudocolored. Quantification of the number of ECs selected to branch into ISVs (C), percentage of ECs that undergo proliferation (E) and total number of ECs per ISV (F) in control, *dll4* knockdown, *flt1* knockdown and SU5416-treated embryos. (G) Illustration of the biphasic nature of EC motile selection in control embryos and upon *dll4* or *flt1* knockdown. (H) Putative role of a VEGFR-regulated bipotential switch in modulation of the temporal dynamics of EC motile selection. (I) Outline of the signaling interactions underpinning construction of the 2-cell ODE mathematical model. Blue arrow indicates lateral inhibition between cells. Green and red arrows indicate positive- and negative-feedback via VEGFR, respectively. HE refers to the combined effects of Notch-induced expression of the transcriptional repressors, HES, HEY and HER (see Methods for more details). (J-K) Plots of DLL4 levels in coupled cells following model simulations using varying levels of positive-feedback. Depending on their final level of DLL4, cells were assigned as having acquired either high VEGFR activity and stable tip identity (high levels DLL4; blue arrowhead; J) or high Notch activity and repressed tip identity (low levels DLL4; red arrowhead; K). Blue and red dashed lines represent max and min DLL4 thresholds for stable tip identity and repressed tip identity, respectively. Increasing levels of positive-feedback consistently reduced the threshold of Vegfr activity required for fast tip cell patterning. Error bars: mean ± SEM. \**P*<0.05, two-way ANOVA test. Scale bars, 25 μm. See also Figure S1.

To define the timing of EC decision-making *in-vivo*, we probed the dynamics of lateral inhibition-mediated motile EC selection during zebrafish intersegmental vessel (ISV) angiogenesis. Live-cell imaging of ISV sprouting in *Tg(kdrl:nlsEGFP)^zf109^* zebrafish embryos revealed that lateral inhibition defines a tight temporally restricted selection window that robustly generated two motile ECs per vessel by 24 hours post fertilisation (hpf; Figure 1B-1C). Importantly, Notch signalling critically closed the window, as in the absence of *dll4* the rate of EC selection did not initially change, but simply continued unstopped (Figure 1C-1D). Unexpectedly, Vegfr levels determined the number of ECs selected within the window, as loss of *flt1* to enhance Vegfr signalling increased the number of ECs selected to sprout, but all were selected within the normal temporal window (Figure 1C). Likewise, the opposite was observed upon brief low-dose inhibition of Vegfr signalling with less cells selected in the time frame (Figure 1C). As we cannot interrogate the signalling dynamics occurring during the selection time window *in vivo*, we utilized the well validated computational Memagent-Spring (MSM) model of Vegf-Notch selection to simulate and predict how cells collectively compete within the DA prior to sprouting[22–24] (see Methods). Using exactly the model parameters as previously published, a single early time window could indeed be found that remarkably exhibited exactly matching phenomena as seen *in vivo* (Figure S1A-S1D), where Vegf-inhibited conditions (vegf=0.038) selected fewer ECs and *fltl* knockdown conditions (Vsink=8) selected more (see methods for full description of these model parameters). Monitoring of signalling dynamics in simulations confirmed that Vegf levels determined the number of selected ECs by modulating the speed of selection within this time window with the model interestingly predicting this is due to the control conditions being suboptimally slow at selecting within this short time window compared to flt1 KD (Fig 1C).

Importantly, differences in Vegfr-dependent selection rates were not associated with differential EC proliferation (Figure. 1E), although a later switch to Notch-dependent mitosis was revealed by *dll4* knockdown. Moreover, proliferation had minimal effect on overall EC numbers, with both the initial rate of Vegfr-dependent EC induction and Notch-dependent closure of the selection window being the primary determinants of vessel cellularity (Figure 1F). Hence, we reveal that temporal control of angiogenesis is distinctly biphasic, involving (1) Vegfr-level-dependent selection of motile EC numbers, then (2) termination of further selection by Dll4-Notch (Figure 1G).

### Positive-feedback temporally regulates lateral inhibition network dynamics

The abrupt switch-like nature of the EC selection window at 24 hpf (Figure. 1C) suggested network ultrasensitivity, whereby a distinct threshold of Vegfr signalling both commits ECs for motile selection and triggers lateral inhibition. Hence, fluctuating Vegfr levels would determine the magnitude of EC selection by dictating how many ECs achieve threshold levels (Figure 1H). Such ultrasensitive responses are frequently driven by positive-feedback loops that amplify signal outputs to promote switch-like dynamics[25,26], but despite the recognized role of Notch-mediated negative-feedback in angiogenesis, the function/identity of positive-feedback modulators remains elusive. Using our previously validated ordinary differential equation (ODE) mathematical model of DLL4-Notch-mediated lateral inhibition[16], which permits rigorous, mathematical interrogation of the bifurcation dynamics in this system, we probed the ability of positive-feedback to modulate the thresholding of EC lateral inhibition. In this model, two adjacent un-patterned ECs compete for selection as either a VEGFR-active DLL4-expressing tip cell or Notch-active laterally inhibited cell using the well-established VEGFR-DLL4-Notch negative-feedback loop (Figure 1I). We created a parameter *P* (see Methods) that creates a positive-feedback interaction whereby VEGFR increases the level/activity of an additional factor that positively feeds back to activate more VEGFR. Using previously defined reaction parameters [16,27–29], we revealed that increasing levels of positive-feedback amplified small differences in VEGFR activity between coupled ECs to drive rapid reciprocal VEGFR/Notch activation, even at low VEGFR levels normally incapable at driving EC sprouting (Figure1J-1K and S1E-S1F). Hence, positive-feedback decreases the threshold of VEGFR activity required to induce motile EC selection and increases the speed of EC decision-making by invoking ultrasensitive switch-like behavior during lateral inhibition.

### Vegfr-induced expression of *tm4sf18* promotes positive-feedback

To support *in silico* observations we expanded upon our previous transcriptomic study [30] to identify putative positive-feedback regulators of Vegfr by defining genes transcriptionally activated by Vegfr and repressed by Notch signaling in zebrafish ECs (Figure 2A). Of only 10 candidate Vegfr/Notch-regulated transcripts we identified *h2.0-like homeobox-1* (*hlx1*), a known transcriptional target of Vegfr activity *in-vivo* [30–32] and the atypical tetraspanin *transmembrane 4 L6 family member 18* (*tm4sf18*). TM4SF family proteins are known membrane-associated adaptors that ligand-independently activate receptor tyrosine kinase activity[33,34]. Moreover, *TM4SF1*, the human homologue of *tm4sf18* (Figure S2), is known to modulate EC motile behavior in-vitro[35,36]. Hence, *tm4sf18* represents an ideal candidate positive-feedback modulator of EC motile identity. The highly dynamic nature Vegfr/Notch-regulated expression of *tm4sf18* was validated via qPCR upon inhibition of Vegfr signaling and *dll4* knockdown (Figures 2B-2C), indicating that *tm4sf18* transcription may be tightly restricted to sprouting EC populations (Figure 2D). This was confirmed following characterization of the spatiotemporal pattern of *tm4sf18* expression during zebrafish development. During early ISV sprouting from 22 to 26 hpf, *tm4sf18* expression was almost exclusively restricted to sprouting ISVs (blue brackets in Figure 2E) and was excluded from adjacent non-angiogenic vascular tissues, such as the dorsal aorta (DA). Importantly, the absence of *tm4sf18* in *cloche (clo^s5^)* mutants that lack endothelial tissues[37] confirmed expression in sprouting ECs (Figure 2F). Moreover, *tm4sf18* expression was ectopically expanded to non-angiogenic tissues upon *dll4* knockdown, demonstrating a tight association with EC sprouting potential (Figure 2F). Indeed, rapid repression of *tm4sf18* was observed following fusion of adjacent ISVs to form the dorsolateral anastomotic vessel (DLAV) and subsequent termination of Vegf-induced angiogenic behavior (red brackets in Figure 2E). Hence, expression of *tm4sf18* is dynamically modulated by the Vegfr-Notch axis and is tightly spatiotemporally restricted to sprouting ECs *in-vivo*.

**Figure 2.**
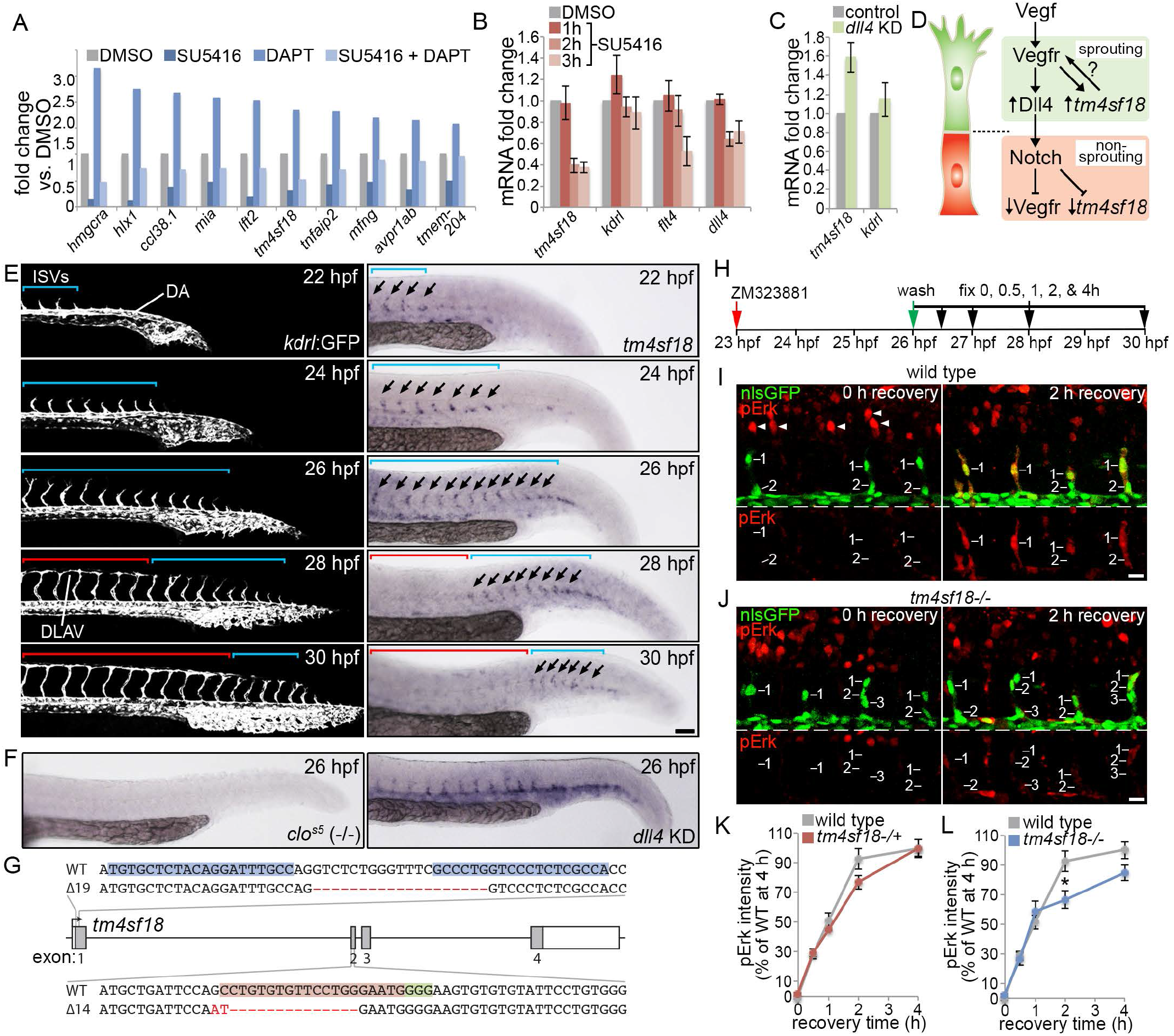
Tm4sf18 is a positive-feedback modulator of Vegfr signaling dynamics. (A) Fold change in transcript levels of the indicated genes following inhibition of Vegfrs (2.5 μM SU5416), Notch (100 μM DAPT) or both from 22 to 30 hpf. GFP-positive ECs were isolated from inhibitor-treated *Tg(kdrl:nlsEGFP)^zf109^* embryos by flow cytometry prior to transcriptome profiling. (B) Fold change in *tm4sf18, kdrl, flt4* and *dll4* transcript levels by qPCR in embryos incubated with SU5416 for the indicated periods of time. SU5416 rapidly down-regulates *tm4sf18* expression. (C) Fold change in *tm4sf18* and *kdrl* transcript levels by qPCR upon *dll4* knockdown. In the absence of Dll4-Notch signaling *tm4sf18* expression was significantly up-regulated. (D) Illustration of proposed regulation of *tm4sf18* expression in sprouting ECs. (E) Lateral views of sprouting intersegmental vessels (ISVs) in *Tg(kdrl:GFP)^s843^* embryos (left panels) or WT embryos following whole-mount *in situ*hybridization analysis of *tm4sf18* expression (right panels) at the indicated developmental stages (DA, dorsal aorta; DLAV, dorsolateral anastomotic vessel; Blue brackets indicate nascent ISVs; Red brackets indicate anastomosed ISVs; arrows indicate *tm4sf18*-expressing ISVs). (F) Whole-mount *in situ* hybridization analysis of *tm4sf18* expression in clo^s5^ mutant embryos or WT embryos upon *dll4* knockdown. (G) Illustration of lesions introduce into the *tm4sf18* genomic loci by TALEN/CRISPR-mediated gene editing. A 19 bp deletion of *tm4sf18* exon 1 and a 16 bp deletion / 2 bp insertion of exon 2 were generated using TALENs and CRISPR/Cas9, respectively. Genomic target sites for the TALENs, gRNA target site and PAM sequence are indicated by blue, red and green highlights, respectively. (H) Strategy for assessing Vegfr signaling dynamics *in-vivo*. Vegfr signaling was first blocked upon incubation of embryos with 0.3 μM ZM323881 for 3 h prior to inhibitor washout for 0, 0.5,1, 2 and 4 h and pErk staining. (I-L) Lateral views of pErk immunostaining in ISV ECs of control (I) or *tm4sf18*^-/-^ homozygous mutant (J) *Tg(kdrl:nlsEGFP)^zf109^* embryos at 0 and 2 h after inhibitor washout and quantification of pErk fluorescence intensity, in *tm4sf18*^+/−^ heterozygous (K) or *tm4sf18*^-/-^ homozygous (L) mutant embryos. Arrowheads in I indicate pErk in neuronal cells. Error bars: mean ± SEM. \**P*<0.05, two-tailed t test. Scale bars, 100 μm (E) 25 μm (I-J). See also Figure S2.

To define *tm4sf18* as a putative temporal modulator of Vegfr/EC signaling *in-vivo*, we used both TALEN and CRISPR/Cas9-mediated gene editing to introduce frame-shift mutations into the long (exon-1) or both long and short (exon-2) isoforms of *tm4fsf18*, respectively (Figure 2G). Importantly, the amplitude of Vegfr signaling in sprouting ECs was unaffected by *tm4sf18* exon-1 and exon-2 mutation (data not shown), indicating that in the absence of Tm4sf18 ECs can still achieve Vegfr activity thresholds sufficient to drive EC motile selection. However, the temporal dynamics of Vegfr signaling were significantly perturbed in homozygous embryos (Figure 2H-2L). To test this, firstly Vegfr activity was fully blocked in ECs of sprouting ISVs that had already emerged from the dorsal aorta upon incubation of embryos for 3 h with the Vegfr inhibitor, ZM323881. Full disruption of Vegfr activity by ZM323881 was confirmed by a lack of immunoreactivity for pErk (Figure 2I-2L), a well-established readout for Vegfr signaling in sprouting ECs [22,38,39]. Moreover, disruption of pErk was specific to ECs, as levels remained unchanged in neighboring neuronal cells (Figures 2I-2J). Inhibitor was then washed out and recovery of EC pErk levels were monitored over the next 4 h. Quantification of pErk in sprouting tip ECs revealed that normal levels were recovered after just 2 h in both control and *tm4sf18*^+/−^ heterozygous mutant embryos (Figure 2I, 2K). However, recovery of Vegfr-mediated pErk stalled after 1 h of recovery and was significantly disrupted in *tm4sf18*^-/-^ homozygous mutants (Figure 2J, 2L). Hence, consistent with a functional role as a positive-feedback modulator of Vegfr activity, expression of *tm4sf18* amplifies EC signaling *in-vivo* and temporally modulates acquisition of high thresholds of Vegfr-dependent pErk.

### Tm4sf18 promotes EC selection and appropriate cellularity of sprouting vessels

To define the functional role of Tm4sf18-mediated positive-feedback in EC decisionmaking *in-vivo* we quantified the rate of EC selection during ISV sprouting to reveal a significant reduction in *tm4sf18*^-/-^ homozygous embryos (Figure 3A). These observations were consistent with model predictions that in the absence of positive-feedback, fewer ECs achieved the Vegfr selection threshold within a defined selection window (Figure 1J-K, S1E-F). In parallel, we noted that resulting hypocellular ISVs in *tm4sf18*^-/-^ homozygous embryos were often shorter (Figure 3B), an observation that was confirmed upon quantification of EC tip (cell 1) and stalk (cell 2) movement (Figure 3C-3D). This phenotype was reliant on mutation of both short and long Tm4sf18 isoforms, as exon1 homozygous mutant embryos were unaffected (Figure S3A), and was not due to indirect differences in ISV morphology (Figure S3B). Importantly, disruption of ISV extension did not appear to be a consequence of reduced EC motility, as movement of emerging *tm4sf18*^-/-^ homozygous ECs was initially indistinguishable from wild type and stalling was only observed later in development (Figure 3C-3D). Strikingly, we observed a near-identical phenotype upon disruption of EC proliferation using hydroxyurea/aphidicolin[39,40] (HU/Ap; Figure 3E), suggesting common underlying defects. Although HU/Ap did not disrupt EC motile selection (Figure S3C; unlike loss of *tm4sf18)* and mutation of *tm4sf18* did not disrupt EC proliferation (Figure 3F; unlike HU/Ap) both *tm4sf18* mutation and HU/Ap did generate very similar reductions in ISV cellularity (Figure 3G-3H). Moreover, quantification of tip EC motility in ISVs containing 1, 2 and 3 or more ECs revealed that this cellularity critically determines vessel extension (Figure 3I). Indeed, consistent with the average number of ECs per ISV in *tm4sf18*^-/-^ mutants (approximately two; Figure 3H), ECs lacking Tm4sf18 on average display the normal extension of cells in hypocellular vessels (Figure. 3I). Likewise, wild type and HU/Ap-treated tip ECs display movements consistent with the average cellularity of vessels in these conditions (Figure S3D). Consequently, both *tm4sf18*^-/-^ mutation and HU/Ap-treatment significantly perturbs the supply of ECs to form the DLAV (Figures 3J-3K). Hence, Tm4sf18 modulates lateral inhibition-dependent selection of ECs that critically determines the cellularity of vessels, which in turn determines the rate of extension of sprouting vessels.

**Figure 3.**
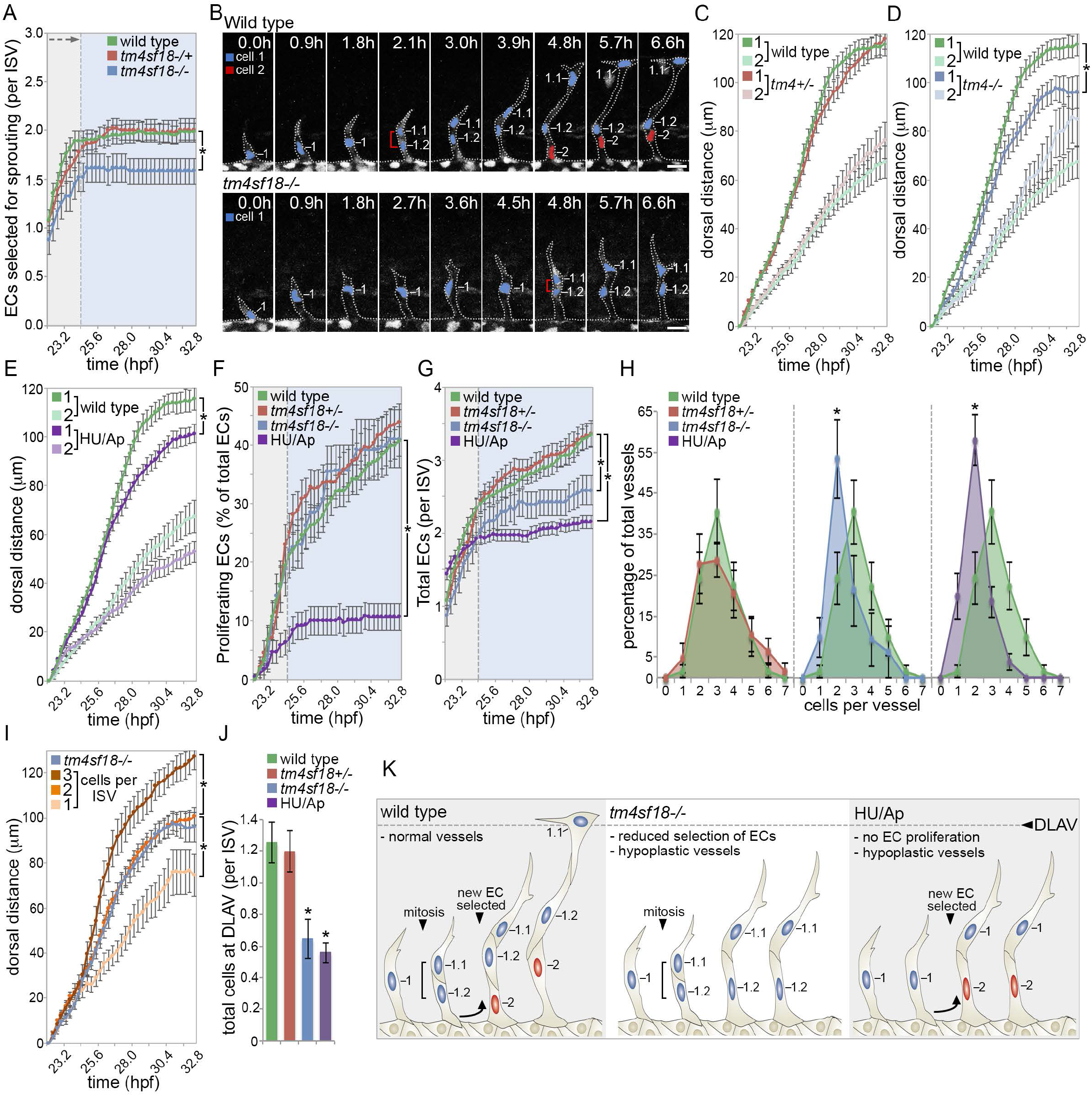
Tm4sf18 regulates lateral inhibition-mediated selection of motile ECs. (A) Quantification of the number of ECs selected to branch into ISVs of wild-type, *tm4sf18*^+/−^ heterozygous and *tm4sf18*^-/-^ homozygous mutant embryos. (B) Time-lapse images of sprouting ISVs in wild type (B) and *tm4sf18*^-/-^ homozygous mutant (C) *Tg(kdrl:nlsEGFP)^zfI09^* embryos from around 19 hpf. Brackets indicate dividing cells. Nuclei are pseudocolored. (C-E) Quantification of the dorsal movement of tip (cell 1) or stalk (cell 2) ECs in wild type, *tm4sf18*^+/−^ heterozygous mutant (C), *tm4sf18*^-/-^ homozygous mutant (D) or 5-hydroxyurea/aphidocolin (HU/Ap)-treated embryos (E). (F-H) Quantification of the percentage of ECs that undergo proliferation (F), total number of ECs per ISV (G) and distribution of ISV cellularity (H) in wild type, *tm4sf18*^+/−^ heterozygous mutant, *tm4sf18*^-/-^ homozygous mutant and HU/Ap-treated embryos. (I) Quantification of the dorsal movement of tip ECs in ISVs consisting of 1, 2 and 3 or more ECs and comparison with the motility of tip ECs in *tm4sf18*^-/-^ homozygous mutant embryos. (J) Quantification of the number of ECs that reach the DLAV position in wild type, *tm4sf18*^+/−^ heterozygous mutant, *tm4sf18*^-/-^ homozygous mutant and HU/Ap-treated embryos. (K) Illustration of the phenotypic consequences of disrupting EC motile selection or proliferation on ISV extension. Error bars: mean ± SEM. \**P*<0.05, two-way ANOVA or two-tailed t test. Scale bars, 25 μm. See also Figure S3.

### Tm4sf18-mediated positive-feedback temporally modulates EC decision-making

Our observations indicated that Tm4sf18 amplifies Vegfr activity to determine the number of ECs that achieve selection threshold. We therefore reasoned that Vegfr inhibition would severely disrupt EC selection in *tm4sf18*^-/-^ homozygous mutants, as ECs would no longer be able to achieve selection threshold in the absence of positive-feedback-mediated amplification of Vegfr signaling. To test this, we blocked EC Vegfr activity with low-dose inhibitor to putatively force ECs to be reliant on positive-feedback amplification of Vegfr activity. Indeed, upon Vegfr inhibition, not only was the emergence of the first selected ECs now greatly delayed in *tm4sf18*^-/-^ homozygous mutants (Figure 4A-4B), this treatment also generated a large delay to the EC selection window (Figure 4B), independent of any effects on EC proliferation (Figure 4C). Hence, when Vegfr activity is limiting, Tm4sf18-mediated positive-feedback is critical to ensure timely EC selection and temporally defines the dynamics of EC lateral inhibition-mediated EC decision-making.

**Figure 4.**
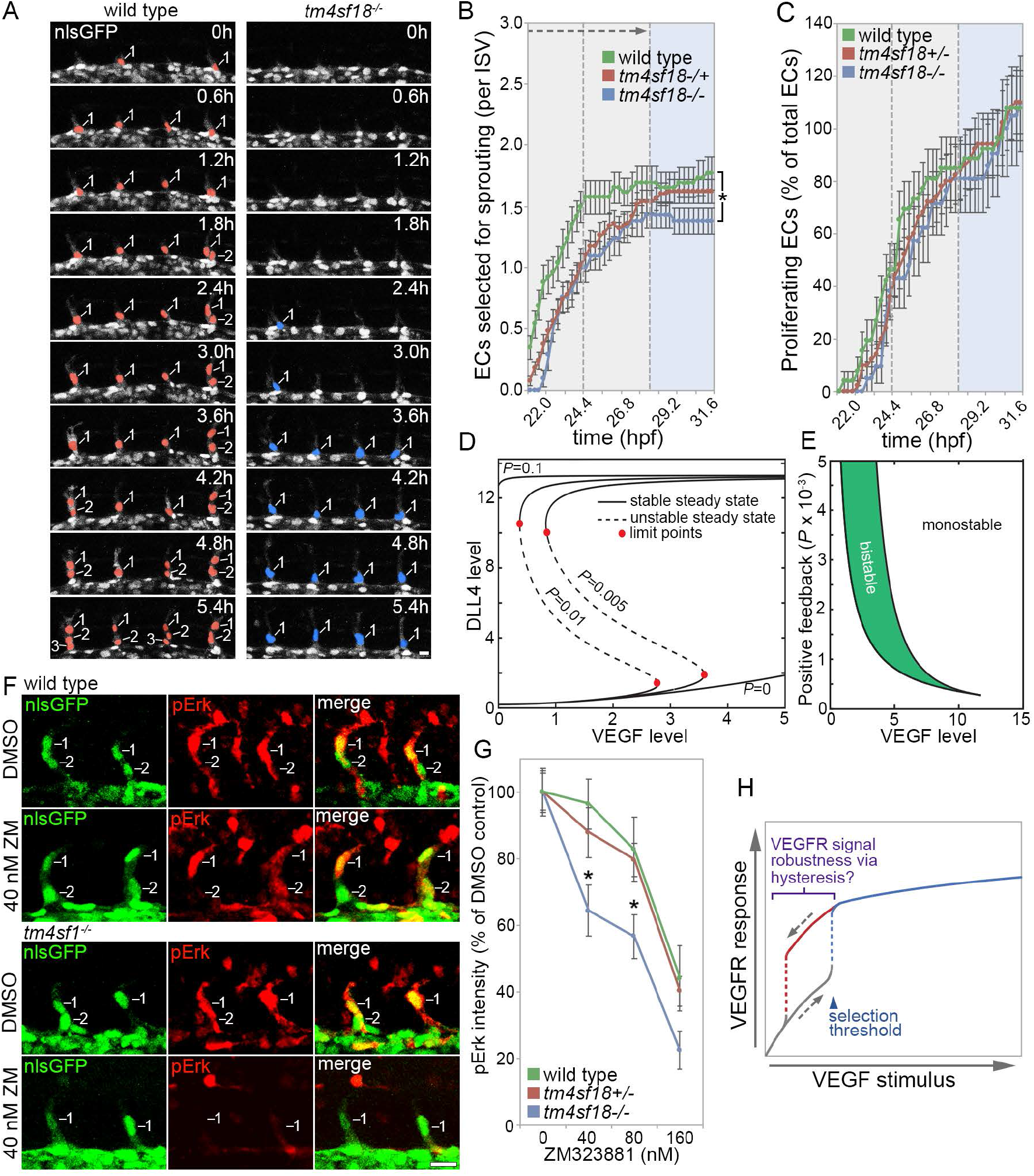
Tm4sf18 defines the timing and robustness of EC decision-making. (A) Time-lapse images of sprouting ISV ECs in either control or *tm4sf18*^-/-^ homozygous mutant *Tg(kdrl:nlsEGFP)^zf109^* embryos from 20 hpf. Embryos were incubated with 40 nM ZM323881 from 18 hpf onward. Nuclei of sprouting ECs emerging from the DA are pseudocolored. (B-C) Quantification of the number of ECs selected to branch (B) and the percentage of ECs that undergo proliferation (C) in 40 nM ZM323881-treated wild type, *tm4sf18*^+/−^ heterozygous mutant and *tm4sf18*^-/-^ homozygous mutant embryos. (D) ODE modelling of the impact of the level of positive-feedback between Tm4sf18 and VEGF *(P)* on network bistability. When there is no positive-feedback, ECs are not able to switch to an active steady state (high DLL4) even when surrounding VEGF is increased. With increasing P, there is a bistable switch in EC state with increasing surrounding VEGF; but at very high *P* values, there is no bistable switch with ECs remaining in an active state with changing VEGF. (E) Two-parameter bifurcation plot with changing VEGF and changing *P* values. Region inside the cusp (i.e. green shaded portion) represents values that are bistable in EC active state with changing values of *P* and VEGF. Everything outside this narrow belt is monostable. (F) Lateral views of sprouting ECs in ISVs of control and *tm4sf18*^-/-^ homozygous mutant *Tg(kdrl:nlsEGFP)^zf109^* embryos immunostained for pErk. Prior to fixation, embryos were incubated with DMSO or 40 nM ZM323881 from 22 hpf for 3 h. (G) Quantification of pErk fluorescence intensity in wild type, *tm4sf18*^+/−^ heterozygous mutant and *tm4sf18*^-/-^ homozygous mutant embryos following incubation with DMSO and increasing concentrations of ZM323881. Upon *tm4sf18* mutation, Vegfr signaling levels were not robust to subtle Vegfr inhibition. (H) Putative role of positive-feedback-generated hysteretic dynamics in the control of VEGFR signal levels and robustness in angiogenesis. Error bars: mean ± SEM. \**P*<0.05, two-way ANOVA or twotailed t test. Scale bars, 25 μm.

### Positive-feedback generates lateral inhibition network robustness

A core feature of positive-feedback is that it contributes to the establishment of bistable networks, which in turn can confer robustness on cell state transitions by employing hysteresis[25,26]. This reinforces robust cell identity decisions by ensuring that lower input signal levels are required to maintain an active cell state than were needed to drive the initial state transition. Further extension of the ODE modeling revealed that increasing levels of positive-feedback (as provided by Tm4sf18) generated such hysteretic dynamics during Vegfr-Notch-mediated lateral inhibition *in silico* (Figure 4D). However, the range of values that generated bistability was narrow (Figure 4E), which may potentially explain why the temporal dynamics of lateral inhibition are also severely disrupted in *tm4sf18*^+/−^ heterozygous mutants upon Vegfr inhibition (Figure 4B). To test these computational predictions that Tm4sf18-mediated positive-feedback could promote robustness of selected EC identity to variation in inductive signal, we waited until after ECs were selected for branching, i.e. already in ISV sprouts (>22 hpf), and hence already above selection thresholds of Vegfr activity. Then we determined the robustness of signaling to increasing concentrations of Vegfr inhibitor for 3h. In wild type and *tm4sf18*^+/−^ heterozygous embryos, Vegfr signaling was highly robust to partial inhibition of Vegfr, with no significant reduction in pErk levels observed upon incubation with 40 nM and 80 nM ZM323881 (Figures 4F-4G). In contrast, Vegfr signaling was no longer protected in *tm4sf18*^-/-^ homozygous mutant embryos and pErk levels were significantly disrupted upon incubation of embryos with low dose inhibitor. Hence, as predicted *in silico* (Figure 4D-4E), Tm4sf18-mediated positive-feedback generates signal robustness reminiscent of a bistable network and buffers Vegfr signaling output to maintain robust angiogenic responses, despite fluctuations to input signal (Figure 4H).

### Conclusions

Using an integrated *in silico* and *in-vivo* approach we provide evidence that Tm4sf18-mediated positive-feedback generates an ultrasensitive switch that directs the timing and robustness of angiogenesis. Recent work demonstrates that such temporal control of EC lateral inhibition ultimately defines the topology of vascular networks, with faster rates of tip EC selection dramatically increasing vessel network density[14]. As such, it will be critical to determine if Tm4sf18-mediated control of Vegfr activity and lateral inhibition underpin such modulation of vessel branch density in more complex vascular beds. Moreover, observations that positive-feedback confers robustness against fluctuations in Vegfr activity indicates that human *TM4SF* genes may be exploited in therapeutic contexts. For example, abnormally high levels of VEGFR signaling were recently shown to drive synchronous oscillations of Dll4 in neighboring EC, underpinning a switch from normal EC communal branching behavior to the pathological vessel expansion associated with human retinopathies[15]. Disruption of positive-feedback would significantly increase the threshold of VEGFR activity required to switch vessels to abnormal expansion, putatively blocking progression to synchronous oscillatory dynamics and defining a novel therapeutic approach to normalize branching. Likewise, disruption of positive-feedback could increase the potency of existing VEGFR-targeting anti-angiogenic anti-cancer therapeutics by reducing the concentration of compound required to block functional angiogenesis. This is especially pertinent considering that a key complication of current anti-angiogenic therapeutics is the dose-limiting side-effects encountered with such drugs[40]. Overall, our observations reveal that the relatively slow dynamics of lateral inhibition-mediated cell fate decisions can be transformed into quick, adaptive and robust decision-making processes by simply incorporating positive-feedback. Excitingly, this study suggests a new general mechanism for temporal adaptation of collective cell fate decisions by positive feedback.

## Acknowledgements

We thank B. Plusa, R. Das, K. Dorey and members of the Herbert lab for helpful discussion and suggestions. We thank members of the University of Manchester Biological Services Facility for providing excellent fish husbandry. S.P.H. is funded by the Wellcome Trust (Ref. 095718/Z/11/Z), BBSRC (Ref. BB/N013174/1) and BHF (PG/16/2/31863). K.B. is funded by BIDMC and NSF (Ref. 1517390) and the Knut and Alice Wallenberg Foundation. T.T. is funded by the Leverhulme Trust (Ref. RPG-2014-370).

## Author contributions

S.P.H. and K.B. conceived the project and designed the experiments. D.J.P. and S.P.H. carried out most of the experiments, while R.T., L.V., T.T. and K.B. carried out some of the experiments. S.P.H., K.B., D.J.P. and R.T. analyzed the data. S.P.H. and K.B. composed the manuscript.

## Declaration of interests

The authors declare no competing interests.

## Methods

### Zebrafish strains and husbandry

Establishment and characterization of the *cloche (clo^s5^)*mutant, *Tg(kdrl:GFP)^s843^* and *Tg(kdrl:nlsEGFP)^zf109^* lines have been described elsewhere [37,41,42]. Embryos and adults were maintained under standard laboratory conditions as described previously [22] and were approved by the University of Manchester Ethical Review Board and performed according to UK Home Office regulations.

### Time-lapse imaging

Confocal microscopy of live *Tg(kdrl:nlsEGFP)^zf109^* embryos was performed as previously [22,30]. Briefly, embryos were mounted in 1% low-melt agarose in glass bottom dishes, which were subsequently filled with media supplemented with 0.0045% 1-Phenyl-2-thiourea and 0.1% tricaine. Embryos were imaged using a 20x dipping objectives on a Zeiss LSM 700 confocal microscope. Embryos were maintained at 28°C and stacks were recorded at every 0.3 h. Tracking of cell motility was performed in ImageJ using the manual tracking plugin. All cell tracking recordings were normalized at each time point relative to the position of the dorsal aorta to account for any dorsal or ventral drift of embryos during imaging.

### Morpholino oligonucleotide (MO) injections

To knock down gene expression, embryos were injected at the one-cell stage with 5 ng control MO, 5 ng *dll4* MO or 3 ng of*flt1* MO. MO sequences were:

5’- CCTCTTACCTCAGTTACAATTTATA -3’ (control)
5’- ATATCGAACATTCTCTTGGTCTTGC – 3’ *(flt1)* [43]
5’- GTTCGAGCTTACCGGCCACCCAAAG -3’ (*dll4*) [6].

All MOs were purchased from Gene Tools.

### Pharmacological treatments

Embryos were manually dechorionated and incubated with compounds from 22 hpf (unless otherwise stated). The following compounds were used in this study: SU5416 (2.5 μM), ZM323881 (40 nM, 80 nM and 160 nM), 5-hydroxyurea (150 μM) and aphidicolin (20 mM).

### Simulations with the memAgent-spring model (MSM)

The MSM model has been well validated against *in vivo* mouse and zebrafish ISV data in previous studies of collective cell dynamics during of Vegf-Notch-mediated tip cell selection, so it made a good choice for simulating the dynamics within the time window observed *in vivo*. In this model, the endothelial cell outer membrane is represented at a subcellular level by a collection of individual computational agents (‘memAgents’) connected by springs following Hooke’s law, which represents the actin cortex beneath. The MSM allows subcellular level rules to generate localized responses of individual memAgents on the cell surface and complex cell shape changes during cell migration.

#### Model initialization and parameterization

The model was initialized with 8 cells in a row, one per vessel cross section (See Fig S1D), representing a collection of endothelial cells in the DA competing to sprout into the ISV space above (represented very simply here as just a fixed vegf gradient extending into the y axis above the horizontal row of cells). All parameters were kept the same as previously published[14,27,44] except those being varied to match the experimental conditions here, described below.

The model was run 100 times for a maximum of 200 timesteps under a range of *vegf* and *flt1* inhibition conditions to see if a single early time window during selection might also generate fewer or more cells being selected by those times as seen *in vivo* for some values of the respective vegf perturbation conditions. VEGF – vegfr activation (*V_m_^′^*) of the vegfr (*V*) in a given memAgent *m* in the model is encapsulated by equation 1 (fully described in[44]) below:

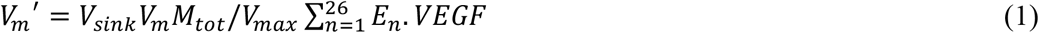

Where, *Vsink* (normally set to 9) is a fixed value which acts as a sink (mimicking *flt1*) reducing the amount of available *VEGF* in the 26 neighbouring environmental grid sites (*E_n_.VEGF*) surrounding that memAgent *m* (as the model runs on the 3D gridded lattice) for binding to its main vegf receptors *V_m_* (only the main receptor V is able to trigger cell migration and Dll4 upregulation in the cells). M_tot_ is the total number of memAgents currently comprising the cell, and V_max_ is the maximum number of receptors the cell can have. This is the only equation that was varied here, by simply reducing the levels of *VEGF* (to model vegf inhibition) and *Vsink*(to model *flt1* loss) respectively.

### ODE model construction & simulation

Interaction between two ECs at the growing end of the angiogenic front have been captured using coupled ordinary differential equations and the 2-cell model previously described [16]. Reactions for the ordinary differential equations of the 2-cell model were written following mass-action kinetics. Details of model construction, list of ODEs, reaction equations and parameters can be found in [16]. Positive-feedback between VEGF and a VEGF-induced/activated factor (P) is captured using the equation;

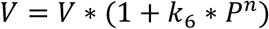

where *k*_6_ is the positive-feedback rate of *P* production and *n* captures the non-linearity of signaling between VEGF (*V*) sensing and *P*. Model simulations were performed using ODE15s solver in MATLAB2013b (www.mathworks.com). All steady state analysis of the ODE model was carried out using the AUTO bifurcation toolbox in XPPAUT (http://www.math.pitt.edu/~bard/xpp/xpp.html).

### Isolation of zebrafish ECs and transcriptome analyses

Flow cytometry-mediated isolation of zebrafish ECs was performed as previously [30]. Briefly, *Tg(kdrl:GFP)^s843^* embryos were dissected and trunks collected in ice cold Ca^2+^/Mg^2+^-free Hank’s buffered salt solution (HBSS), washed four times in 1 ml ice cold Ca^2+^/Mg^2+^-free HBSS and dissociated in 2 ml TrypLE (Invitrogen) at 27.5° C for 30 min with regular agitation. Dissociation was inactivated upon addition of 100 μl fetal bovine serum (FBS). Dissociated cells were subsequently isolated by centrifugation, re-suspended in 5 ml Ca^2+^/Mg^2+^-containing HBSS (with 5% FBS) and passed through 40 μm filters. ECs were collected upon re-centrifugation of dissociated cells, re-suspension in 0.5 ml Ca^2+^/Mg^2+^-containing HBSS (with 5% FBS) and FACS isolation of the kdrl:GFP-positive cell population directly into lysis buffer. Total RNA was isolated using the RNAqueous-Micro kit (Ambion). Complementary DNAs were amplified, labeled with Cy3 (from DMSO-treated embryos) or Cy5 (chemical-treated embryos) and hybridized to the Agilent Zebrafish Gene Expression Microarray (V2) by Mogene Lc. The extracted data were normalized and quality controlled using GeneSpring GX software (Agilent). All probes with a green processed signal below 100 were considered as background.

### Quantitative real-time PCR (qPCR)

Zebrafish embryo or HUVEC cDNA samples were diluted to a concentration of 50 ng/μl with dH_2_0. Each qPCR reaction was prepared in triplicate in a 48 or 96-well plate with each well consisting of 0.2 μM each forward and reverse primer, 50ng cDNA and SYBR Green Mastermix (Applied Biosystems). Reactions were run on an Eco Real-Time PCR System (Illumina) or Step One Plus Real-time PCR System (Applied Biosystems) alongside negative controls. qPCR data was analyzed by the ΔΔC_T_ method and expression normalized to *β-actin* and *ef1a* (zebrafish) or *GAPDH* (human). A relative quantification of gene expression was then determined using the formula 2’^ΔΔCT^.

Primers used for qPCR amplification were:

zebrafish *β actin* forward: 5’-CGAGCTGTCTTCCCATCCA-3’

zebrafish *β actin* reverse: 5’-TCACCAACGTAGCTGTCTTTCTG-3’ [45]

zebrafish *dll4* forward: 5’-TGGCCAGTTATCCTGTCTCC-3’

zebrafish *dll4* reverse: 5’-CTCACTGCATCCCTCCAGAC-3’ [46]

zebrafish *ef1α* forward: 5’-CTGGAGGCCAGCTCAAACAT-3’

zebrafish *ef1α* reverse: 5’- ATCAAGAAGAGTAGTACCGCTAGCATTAC-3’ [45]

zebrafish *flt4* forward: 5’-CTGTCGGATTTGGATTGGGA-3’

zebrafish *flt4* reverse: 5’-GGTGGACTCATAGAAAACCCATTC-3’ [47]

zebrafish *kdrl* forward: 5’ -ACTTTGAGT GGGAGTTTCAT AAGGA-3’

zebrafish *kdrl* reverse: 5’-TTGGACCGGTGTGGTGCTA-3’ [47]

zebrafish *tm4sf18* forward: 5’-CTGGATACTGCTTCCTGATCTC-3’

zebrafish *tm4sf18* reverse: 5’-CAAACAGATACCGTCCCTCAT-3’

### Cloning of *tm4sf18* and whole-mount *in situ* hybridization

The zebrafish *tm4sf18 in-situ* hybridization construct was generated by PCR amplification of the *tm4sf18* ORF from cDNA and cloning of this fragment into pCR-Blunt II-TOPO (Invitrogen). For probe generation, pCR-Blunt II-TOPO *tm4sf18* was linearized with EcoRV at 37°C for 3 h and T7 was used for transcription. For whole-mount *in situ* hybridization, embryos were fixed in 4% paraformaldehyde (PFA) overnight at 4°C and processed as described previously [48].

### Phylogenetic analysis

For phylogenetic analysis of the TM4SF1/4/18 protein family, the NCBI Reference Sequences of TM4SF1, TM4SF4, TM4SF18 proteins of each species were used respectively: human (*Homo sapiens*; NP_055035.1, NP_004608.1, NP_620141.1), mouse (*Mus musculus*; NP_032562.1, NP_663514.2), chicken (*Gallus gallus*; NP_001264407.1, XP_001234023.1, XP_001234069.1), turtle (*Pelodiscus sinensis*; XP_006115992.1, XP_006115993.1, XP_006115991.1), frog (*Xenopus tropicalis*; XP_002937372.1, NP_988958.1, XP_002937374.1), medaka (*Oryzias latipes*; XP_004068091.1, XP_004068090.1), zebrafish (*Danio rerio*; NP_001003489.1, NP_001038487.2), fugu (*Takifugu rubripes*; XP_003974429.1, UniProt H2VCB8), tetraodon (*Tetraodon nigroviridis*; CAF90631.1, CAF90632.1) and elephant shark (*Callorhinchus milii*; XP_007900629.1). Human TM4SF5 protein (NP_003954.2) was used as an out-group for our phylogenetic analysis.

The TM4SF1/4/18 amino acid sequences from all 10 species were aligned using Clustal X2 [49]. The sequences were then manually trimmed of all sites that were not unambiguously aligned. Phylogenetic analysis of amino acid sequences was first performed using NL method implemented in ClustalX2, with outputs displayed using TreeView [50]. Confidence in the phylogeny was assessed by bootstrap re-sampling of the data. For ML tree, the JTT model of protein evolution was used in RAxML [51] with the proportion of invariable sites and gamma parameter estimated from the data, four categories of between-site rate variation; 100 bootstraps were used in the primary ML tree (final ML optimization likelihood: -4945.841564).

### Gene editing

TALENs were designed and constructed to target exon 1 of zebrafish *tm4sf18* using online tools (https://tale-nt.cac.cornell.edu/node/add/talen) and as previously described using the Golden Gate method [52]. The target sequences chosen for the forward and reverse TALENs were 5’-TGTGCTCTACAGGATTTGCC-3’ and 5’-GCCCTGGTCCCTCTCGCCA-3’, respectively. 100 pg of both forward and reverse TALEN mRNA was co-injected into the single cell of zebrafish embryos. At around 24-72 hpf, genomic DNA was extracted from individual embryos and somatic lesions confirmed by high resolution melt (HRM) using Meltdoctor HRM Mastermix (Applied Biosystems) and the following primers:

*tm4sf18* TALEN forward 5’-CTGTTTTCTCCCCCACACAC-3’*tm4sf18* TALEN reverse 5’-TACTCACAGCCAGACCACCA-3’

The CRISPR target site within exon 2 of *tm4sf18* (5’-CCTGTGTGTTCCTGGGAATG-3’) was identified as previously[53]. gRNA and *nls-zCas9-nls* RNA were generated as previously [54]. 30-100 ng of gRNA and 100-150 ng of *nls-zCas9-nls* RNA were co-injected into single cell stage embryos mixed with a phenol red tracer. At around 24-72 hpf, genomic DNA was extracted from individual embryos and somatic lesions confirmed by HRM (as above) using the following primers:

*tm4sf18* CRISPR forward 5’- CATCAGTCTTTGCAGCGAGA -3’

*tm4sf18* CRISPR reverse 5’- TGTAGCATATCCCAACACTCAC -3’

### pErk immunostaining

Whole-mount immunostaining for pErk was performed as previously [22]. Briefly, *Tg(kdrl:nlsEGFP)^zf109^* embryos were fixed in PFA overnight prior to washing in 100% MeOH, incubation with 3% H_2_0_2_ in MeOH on ice for 60 min and further 100%MeOH washes. Embryos were then stored at -20°C for 2 days in MeOH before equilibration with PBT (PBS, 0.1% Tween-20) washes and cryoprotected in 30% sucrose in PBT overnight at 4°C. The next day embryos were equilibrated in PBT, incubated with 150 mM Tris-HCl (pH 9.0) for 5 min and then heated to 70°C for 15 min. Embryos were then washed with PBT and then twice with dH_2_0 for 5 min. Water was then removed prior to addition of ice-cold acetone for 20 min at -20°C. Acetone was removed prior to PBT washes, one TBST (TBS, 0.1% Tween-20, 0.1% Triton X-100) wash and incubation overnight at 4°C with block solution (TBST, 1% BSA, 10% goat serum). The next day embryos were then incubated with anti-phospho-ERK1/2 antibody (1:250, Cell Signaling; #4370) in blocking buffer overnight at 4°C. Washes in TBST at room temperature were followed by a wash in Maleic buffer (150 mM Maleic acid, 100 mM NaCl, 0.001% Tween-20, pH 7.4) for 30 min. Embryos were then blocked in 2% blocking reagent (Roche) in Maleic buffer for 3 h at room temperature prior to incubation with goat anti-rabbit IgG-HRP (1:1000) in 2% blocking reagent in Maleic buffer overnight at 4°C. Embryos were then washed in Maleic buffer and then PBS at room temperature prior to incubation with 50 μl amplification diluent with 1 μl Tyramide-Cy3 (Perkin Elmer) for 3 h at room temperature in the dark. Embryos were finally washed over several days in TBST at room temperature. Levels of pErk were quantified as the mean nuclear Cy3 fluorescence intensity using ImageJ.

### Statistics

All statistical analyses were performed using GraphPad Prism 6.0 or Microsoft Excel software. Statistical significance was assessed using either unpaired two-tailed Student’s *t*-tests or two-way ANOVA tests. Results are presented as mean ± S.E.M. All experiments were repeated at least three times. For all analyses *P*<0.05 was considered statistically significant.

## Supplemental Information

**Figure S1. VEGFR signaling and positive-feedback define the magnitude and timing of tip EC selection in vivo, related to Figure 1.** (A-B) MSM modeling of the influence of differential Vegfr (A) and flt1 (B) levels on the timing of EC activation via lateral inhibition. By defining a selection window at approximately 175-time steps, the model accurately recapitulates the in-vivo behavior induced by Vegfr inhibition (decreased EC selection) and flt1 knockdown (increased EC selection) in Figure 1C. (C) Quantification of the number of cells selected in the selection window both in silico and in-vivo. (D) Example frames from one simulation run of a control simulation (Vsink=9) vs a flt1 KD simulation (Vsink =8) showing by the end of the time window the control is actually less efficient, i.e. slower to select within the time window as the flt1 KD selects all possible tip cells from the available pool of cells more rapidly. Frames taken at t=0,50,100,150,200. (E-F) Matrix plots of EC patterning speeds in the 2-cell ODE system following exposure of each cell to different VEGF levels in the absence (E) or presence (F) of positive-feedback. Grey boxes indicate either the failure to adequately pattern into Vegfr active and Notch active ECs or very slow patterning. Larger orange boxes indicate coupled ECs experiencing the lowest VEGF levels (<0.05 c.u.). Positive-feedback globally increases the speed of cell patterning and greatly reduces the threshold of VEGF levels capable of generating robust patterning (see large orange boxes).

**Figure S2. Phylogenetic, synteny and protein analysis of the TM4SF family, related to Figure 2.** (A-B) Phylogenetic trees of vertebrate TM4SF1/4/18 protein family based on either neighbor-joining (NJ, A) or maximum-likelihood (ML, B) analyses. Human TM4SF5 protein sequence was defined as the out-group. Branch lengths are proportional to evolutionary distance corrected for multiple substitutions; the scale bar denotes 0.1 underlying amino acid substitutions per site. Figures on branches indicate robustness of each node (>50%), estimated from 1000 bootstrap replicates for NJ (A) and 100 replicates for ML (B). All nodes with a bootstrap support of less than 50% were collapsed to a polytomy. Based on separate analyses of all vertebrate TM4SF protein (data not shown), TM4SF1, 4 and 18 form a monophyletic group and likely share a common ancestor. Hence, TM4SF1 and TM4SF18 are the most related in the TM4SF family. (C) Synteny analysis of vertebrate Tm4SF1/4/18 in the human, mouse, chick, turtle and zebrafish genome assemblies. Dashed lines represent breaks in synteny. TM4SF1, 4 and 18 are closely located on the same chromosome, suggesting that they were generated by two tandem gene duplications at early stages of vertebrate evolution. The first of these gave rise to the TM4SF4 and TM4SF1/18 paralogous groups, and the second gave rise to the TM4SF1 and TM4SF18 paralogous groups. The tm4sf1 subfamily appears to have subsequently been lost in the lineage leading to teleost fish. Similarly, the Tm4sf18 subfamily appears to have subsequently been lost in mice. (D) Percentage BLAST sequence identity (black numbers) and sequence similarity (red numbers) between teleost fish (zebrafish, medaka, tetraodon and fugu) Tm4sf18 and human, mouse or chick TM4SF1/18. Following the loss of tm4sf1, the remaining Tm4sf18 in teleost fish consistently shares higher protein sequence identity/similarity with human and chicken TM4SF1 over TM4SF18. Hence, zebrafish Tm4sf18 appears to have acquired the function of Tm4sf1 and is the predominant teleost homologue of mammalian TM4SF1.

**Figure S3. Phenotypic consequences of tm4sf18 mutation and disruption of EC proliferation, related to Figure 3.** (A) Quantification of the dorsal movement of tip ECs in wild type, tm4sf18+/− exon1 heterozygous mutant and tm4sf18-/- exon-1 homozygous mutant embryos. Mutation of only the long isoform of tm4sf18 does not disrupt EC motility. (B) Quantification of the length of any fully formed ISVs observed in wild type, tm4sf18+/− heterozygous mutant, tm4sf18-/- homozygous mutant and hydroxyurea / aphidocolin (HU/Ap)-treated embryos. None of these perturbations affects the morphology of successfully formed ISVs. (C) Quantification of the number of ECs selected to branch into ISVs in wild type or HU/Ap-treated embryos. Disruption of EC proliferation has no significant effect on selection of motile ECs by lateral inhibition. (D) Quantification of the dorsal movement of tip ECs in ISV consisting of 1, 2 and 3 and more ECs and comparison with the motility of tip ECs in wild type and HU/Ap-treated embryos. Tip ECs in wild type embryos behave like those in ISVs with three cells whereas inhibition of proliferation generates tip ECs that behave like those in ISVs with only two cells, consistent with the average number of ECs per ISV in both of these situations (see Figure 3H). Error bars: mean ± SEM.

